# BITE: an R package for biodiversity analyses

**DOI:** 10.1101/181610

**Authors:** Marco Milanesi, Stefano Capomaccio, Elia Vajana, Lorenzo Bomba, José Fernando Garcia, Paolo Ajmone-Marsan, Licia Colli

## Abstract

Nowadays, molecular data analyses for biodiversity studies often require advanced bioinformatics skills, preventing many life scientists from analyzing their own data autonomously. BITE R package provides complete and user-friendly functions to handle SNP data and third-party software results (i.e. Admixture, TreeMix), facilitating their visualization, interpretation and use. Furthermore, BITE implements additional useful procedures, such as representative sampling and bootstrap for TreeMix, filling the gap in existing biodiversity data analysis tools.

**Availability:** <u>https://github.com/marcomilanesi/BITE</u>

## 1 Introduction

In the last decade, thanks to the development of high-throughput sequencing and genotyping devices, the molecular datasets used in genomics reached the “big data” size (Stephens *et al.*, 2015). To face the challenge of big-data handling and analysis, several life scientists have switched to computational biology or bioinformatics (Loman and Watson, 2013). This transition is far from being painless due to a number of factors, as the complexity of the tasks, the time required to acquire programming skills, the difficulty of selecting the appropriate tool among a jungle of available software and, often, the necessity to adapt and integrate multiple analyses into a single pipeline to obtain the desired results.

Here, we introduce BITE (BioInformatics Tools for Everyone), an R package providing optimized and user-friendly functions to analyze single nucleotide polymorphism (SNP) data for biodiversity studies and handle third-party software results (e.g. Admixture, TreeMix), facilitating their visualization and interpretation. Furthermore, BITE returns publication-grade plots, summary statistics and files ready-to-use in third-party software.

## 2 Implementation

### 2.1 Genomic data handling

BITE accepts data in transposed PLINK format (Purcell *et al.*, 2007; Chang *et al.*, 2015) and makes extensive use of GenABEL R package functions (Aulchenko *et al.*, 2007). *bite.qc* is designed to run quality control procedures, simplifying (i.e. use of a single function for the entire process) and improving (i.e. output files with retained/excluded SNP/ID in PLINK format) the *check.marker* function from GenABEL.

Since unequal sample size among populations can bias biodiversity analyses results (e.g. Subramanian, 2016), a frequently adopted strategy is to harmonize sample sizes by randomly subsampling the individuals within large populations. However, whether a given subset of individuals is representative or not of the original population is an issue highly debated within the scientific community, with no universally accepted solution. *representative.sample* function tries to identify a subset of individuals of user-defined size mirroring as closely as possible the genetic structure of the original dataset. First, a Multi-Dimensional Scaling analysis is performed on the matrices of pairwise Identity-by-State distances, from both the original and the subsampled datasets. Then, for each of the *p* dimensions, whose number is user-defined, the distributions of the scores obtained for the original and subsampled sets are compared using a two-sample Kolmogorov-Smirnov test (Young, 1977). If the distributions on the *p* dimensions are not significantly different, then the subset is considered representative of the original population.

Finally, *bite.biostat* function provides summary statistics on markers and individuals, and their graphical representations.

### 2.2 Graphical representation of membership coefficients

To date, several software have been devised to estimate membership coefficients and assign individuals to latent clusters (e.g. Wollstein and Lao, 2015). BITE accepts output files from the most commonly used model-based ancestry estimation software, such as Admixture (Alexander *et al.*, 2009), FastSTRUCTURE (Raj *et al.*, 2014) and sNMF (Frichot *et al.*, 2014), and offers an improved graphical representation, by keeping the color of a given genomic component constant throughout cluster solutions, thus easing the interpretation of results.

Bar plot representations can be returned either in classical fashion using *membercoef.plot* function, or in circular fashion (both full and half circle) with *membercoef.circos* function, as in Circos software (<u>http://www.circos.ca/</u>), using a modified code from RCircos package (Zhang *et al.*, 2013). The circular plot comes in handy particularly when many populations and Ks are plotted at the same time (Figure 1A).

**Fig. 1.**
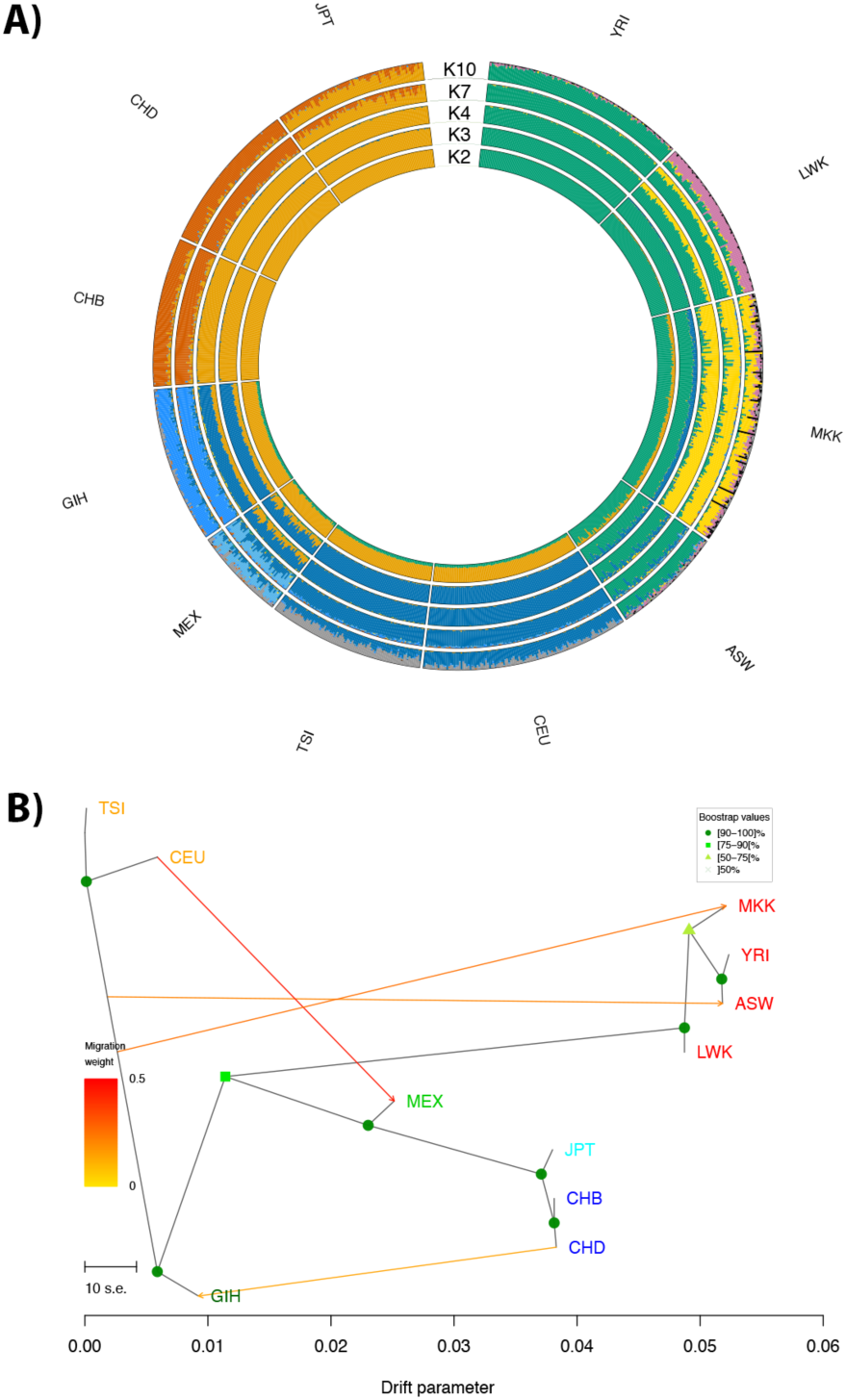
Examples of BITE function outputs from human trial datasets. A) Membership coefficients estimated with Admixture software at K= 2, 3, 4, 7 and plotted in curcular fasihion.with the *membercoef.circos* function of BITE; B) Gene flow analysis using TreeMix. Nodes robustness was estimated with 100 bootstrap replicates and plotted using the *treemix.bootstrap* function of BITE.

Finally *membercoef.cv* helps the user identifying the most likely K by plotting the values of the relevant metrics.

### 2.3 Gene flow analyses

TreeMix software (Pickrell and Pritchard, 2012) infers the patterns of population splits and mixtures. The *treemix.scripts* function provides BASH scripts to run TreeMix software either in a conventional way or with the bootstrap procedure, newly implemented in BITE. Both BASH scripts make use of GNU parallel (Tange, 2011) to exploit multi-thread machines. By estimating the robustness of the tree nodes, the bootstrap procedure strengthens the identification of statistically supported population splits and gene flow events. Briefly, after the number of migrations to test (*m*_*best*_) has been identified, *n* bootstrap replicates of *m*_*best*_ tree are run and used to build a consensus tree with PHYLIP (Felsenstein, 2016). The consensus tree is then loaded in TreeMix to re-estimate the *m*_*best*_ migrations. The result from *Treemix_boostrap.sh* script is plotted using *treemix.bootstrap* (Figure 1B).

The function *newick.split,* used internally, identifies the branch splits within the Newick tree and assigns the respective bootstrap values to the nodes. *treemix.fit* returns the fraction of explained variance in relatedness between populations useful to evaluate the fit of the tested TreeMix models. Genetic drift between populations can be visualized as a heatmap using *treemix.drift*. BITE also includes all original R functions developed by Pickrell and Pritchard (2012) for the graphical representation of TreeMix results.

## 3 Example

BITE functionalities can be tested using a subset of the data from HapMap Project Phase 3 (Altshuler *et al.*, 2010), comprising 1,011 subjects from 11 human populations and 20,000 randomly selected SNPs mapping to chromosome 2. Data examples, available through *bite.trialdata*, include i) binary PLINK files, ii) output files from the aforementioned membership estimation software (for K=2 to K=10) and iii) output from TreeMix software run both in conventional (*m*=0-5) and bootstrap (*m*=4) mode.

## 4 Conclusion

The BITE package provides functions for a wide range of applications in biodiversity studies, allowing users to perform fine tuned, customized and advanced analyses. The package is available at <u>https://github.com/marcomilanesi/BITE</u>.

## Acknowledgements

MM was supported by grant 2016/05787-7, São Paulo Research Foundation (FAPESP). EV was supported by the Doctoral School on the Agro-Food System (Agrisystem) of the Università Cattolica del Sacro Cuore (Italy).

*Conflict of Interest*: none declared.

## References

Alexander, D.H. et al. (2009) Fast model-based estimation of ancestry in unrelated individuals. Genome Res., 19, 1655–1664.

Altshuler, D.M. et al. (2010) Integrating common and rare genetic variation in diverse human populations. Nature, 467, 52–58.

Aulchenko, Y.S. et al. (2007) GenABEL: an R library for genome-wide association analysis. Bioinformatics, 23, 1294–1296.

Chang, C.C. et al. (2015) Second-generation PLINK: rising to the challenge of larger and richer datasets. GigaScience, 4, 7.

Felsenstein J (2016) PHYLIP (PHYLogeny Inference Package) Department of Genome Sciences, University of Washington, Seattle.

Frichot, E. et al. (2014) Fast and Efficient Estimation of Individual Ancestry Coefficients. Genetics, 196, 973–983.

Loman, N. and Watson, M. (2013) So you want to be a computational biologist? Nat. Biotechnol., 31, 996–998.

Pickrell, J.K. and Pritchard, J.K. (2012) Inference of Population Splits and Mixtures from Genome-Wide Allele Frequency Data. PLOS Genet., 8, e1002967.

Purcell, S. et al. (2007) PLINK: A Tool Set for Whole-Genome Association and Population-Based Linkage Analyses. Am. J. Hum. Genet., 81, 559–575.

Raj, A. et al. (2014) fastSTRUCTURE: variational inference of population structure in large SNP data sets. Genetics, 197, 573–580

Stephens, Z.D. et al. (2015) Big Data: Astronomical or Genomical? PLOS Biol., 13, e1002195.

Subramanian, S. (2016) The effects of sample size on population genomic analyses – implications for the tests of neutrality. BMC Genomics, 17.

Tange, O. (2011) GNU Parallel - The Command-Line Power Tool. Login USENIX Mag., 36, 42–47.

Wollstein, A. and Lao, O. (2015) Detecting individual ancestry in the human genome. Investig. Genet., 6, 7.

Young, I.T. (1977) Proof without prejudice: use of the Kolmogorov-Smirnov test for the analysis of histograms from flow systems and other sources. J. Histochem. Cytochem. Off. J. Histochem. Soc., 25, 935–941.

Zhang, H. et al. (2013) RCircos: an R package for Circos 2D track plots. BMC Bioinformatics, 14, 244.

